# Striatal dopamine integrates cost, benefit and motivation

**DOI:** 10.1101/2022.09.20.508777

**Authors:** Neir Eshel, Gavin C. Touponse, Allan R. Wang, Amber K. Osterman, Amei N. Shank, Alexandra M. Groome, Lara Taniguchi, Daniel F. Cardozo Pinto, Jason Tucciarone, Brandon S. Bentzley, Robert C. Malenka

## Abstract

Dopamine (DA) release in the ventral and dorsal striatum has been linked to reward processing and motivation, but there are longstanding controversies about whether DA release in these key target structures primarily reflects costs or benefits, and how these signals vary with motivation. Here we apply behavioral economic principles to generate demand curves for rewards while directly measuring DA release in the nucleus accumbens (NAc) and dorsolateral striatum (DLS) via a genetically-encoded sensor. By independently varying costs and benefits, we reveal that DA release in both structures incorporates reward magnitude and sunk cost. Surprisingly, motivation was inversely correlated with reward-evoked DA release; the higher the motivation for rewards the lower the reward-evoked DA release. These relationships between DA release, cost and motivation remained identical when we used optogenetic activation of striatal DA inputs as a reward. Our results reconcile previous disparate findings by demonstrating that during operant tasks, striatal DA release simultaneously encodes cost, benefit and motivation but in distinct manners over different time scales.

## INTRODUCTION

Decision making requires a consideration of both costs and benefits, as influenced by motivation. Although mesolimbic DA release is crucial for reward learning and therefore decision making^1–4^, prior studies have disagreed over its role in encoding costs, benefits or motivation. On the one hand, disruptions to DA signaling in the NAc with pharmacological manipulations or lesions bias animals towards low-cost, less-preferred rewards rather than high-cost, more-preferred rewards without affecting reward preference or overall consumption^5–8^. These results led to the conclusion that NAc DA release primarily mediates cost calculations; that is, how much effort an animal exerts to obtain rewards. However, these studies often confounded cost and benefit by varying both simultaneously, and used tools that could not examine how DA activity dynamics in specific target structures underlie these behaviors. In contrast, studies of DA activity dynamics during instrumental tasks suggest that striatal DA more reliably encodes prospective benefit, not cost^9–14^. These studies, however, were mostly correlational and did not exploit causal tools such as optogenetics to explore how DA neuron activity directly influences behavior. Furthermore, DA recording studies have rarely explored changes in motivational state between individuals or across time except in studies of addiction, where there is longstanding debate over whether striatal DA release is sensitized^15^ or depleted^16^ once individuals transition to a highly-motivated, addicted state.

To address these limitations of previous work, we established a simple operant task that independently varies costs and benefits to generate behavioral economic demand curves in response to sucrose rewards or optogenetic stimulation of DA inputs. This approach yields a quantitative metric of motivation, defined by the willingness to overcome cost, independently from reward dose or preferred consumption at no cost. At the same time, we measured striatal DA release using a genetically-encoded sensor, allowing us to test how striatal DA release reflects costs, benefits and motivational state. We find that striatal DA release integrates cost and benefit on a trial-by-trial basis, and surprisingly, that high motivation dampens these signals. These findings reconcile discrepancies between prior studies on DA and help clarify the role of striatal DA signals in motivated behavior.

### DA release in NAc and DLS signals both cost and benefit

Mice learned to poke in an active nosepoke hole for access to sucrose reward, accompanied by a brief light-sound cue (Fig. 1a). Over training, mice increased the number of rewards earned in each session and decreased their latency to consume each reward (Fig. 1b). Once they reached a behavioral criterion (see Methods), mice advanced to economic demand sessions, in which we varied the fixed ratio (FR, number of pokes required per reward) in 10-minute bins over 50 minutes (Fig. 1a). We considered the FR as the “cost” of reward. For each session, reward “benefit” was fixed at a single combination of sucrose concentration and quantity. Between sessions, however, we varied the concentration and quantity, such that mice experienced four different benefits. As expected, mouse operant behavior was sensitive to both reward benefit (Fig. 1c, left) and cost (Fig. 1c, right).

**Fig. 1:**
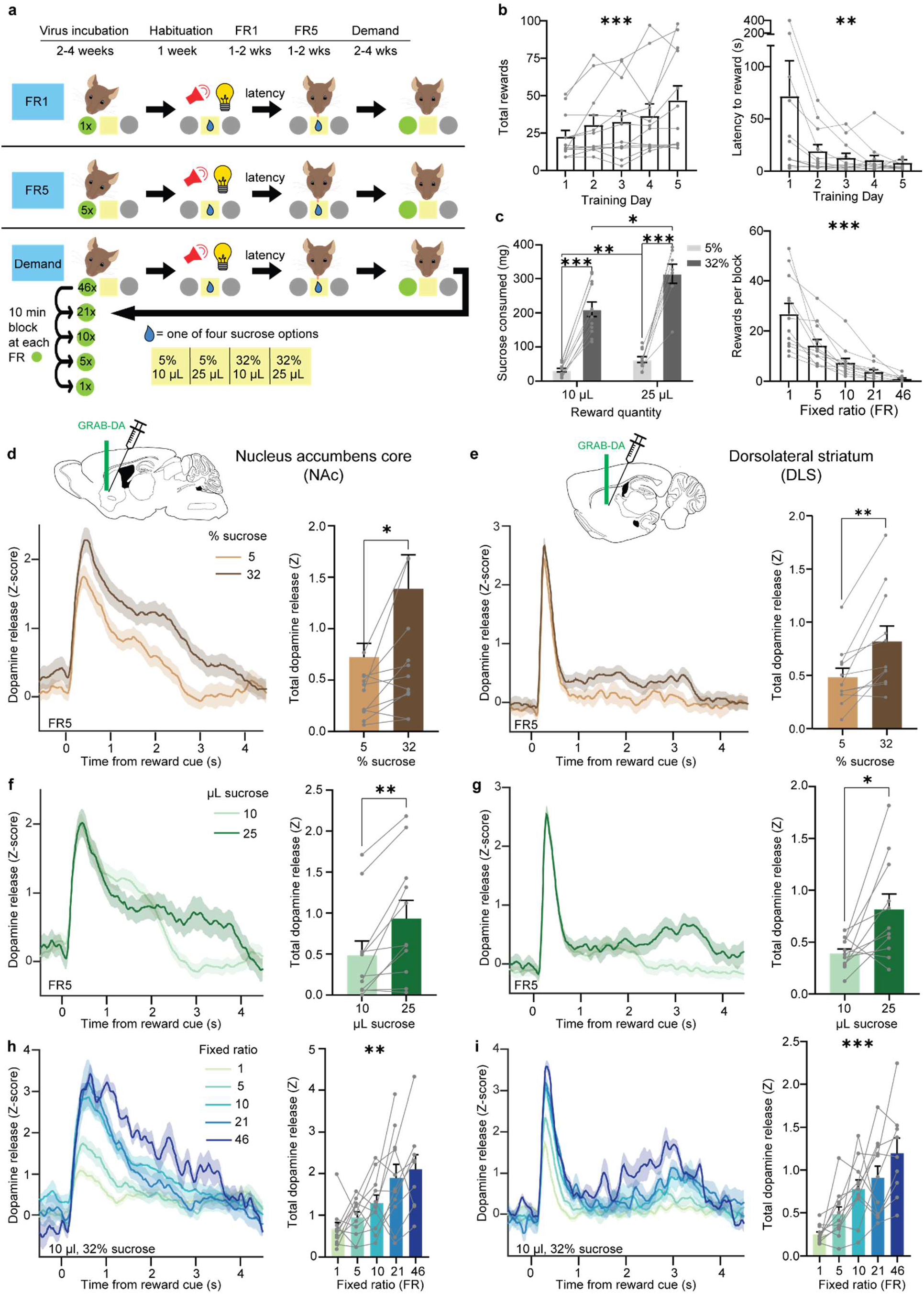
DA release in NAc and DLS encodes both benefit and cost. **a**, Experimental timeline and schematic of behavioral task. FR, fixed ratio. **b**, Left, number of rewards earned each day during FR1 training (Friedman test, Friedman statistic=22.4, P=0.0002; n=12). Right, latency to enter reward port after light-sound cue (Friedman test, Friedman statistic=16.3, P=0.0026; n=12). **c**, Left, sucrose consumed per session as a function of the quantity and concentration of each reward (Mixed-effects model, fixed effect of quantity, F_1,11_=17.9, P=0.0014; fixed effect of concentration, F_1,11_=176, P<0.0001; Tukey-corrected multiple comparisons: 10μL-5% vs 10μL-32%, P<0.0001; 10μL-5% vs 25μL-5%, P=0.0029; 10μL-32% vs 25μL-32%, P=0.024; 25μL-5% vs 25μL-32%, P=0.0001; n=12). Right, average number of rewards earned for each FR (Friedman test, Friedman statistic=47.6, P<0.0001; n=12). **d-i**, Left, Z-scored photometry traces of GRAB-DA fluorescence aligned to reward-predicting cues. Right, area under the curve (AUC) of Z-scored photometry traces from 0 to 4s after cue onset as a function of sucrose concentration (**d**: Wilcoxon matched-pairs signed rank test, W=50, P=0.024; n=12; **e**: Wilcoxon matched-pairs signed rank test, W=64, P=0.002; n=12), sucrose quantity (**f**: Wilcoxon matched-pairs signed rank test, W=62, P=0.003; n=12, **g**: Wilcoxon matched-pairs signed rank test, W=54, P=0.014; n=12) or FR (**h**: Friedman test, Friedman statistic=15.2, P=0.0043; n=12, **i**: Friedman test, Friedman statistic=24.2, P<0.0001; n=12). **d, f, h,** recordings in the NAc. **e, g, i,** recordings in the DLS. *P<0.05, **P<0.01, ***P<0.001. Error bars denote s.e.m.

While mice performed this task, we used fiber photometry and the fluorescent sensor GRAB-DA^17^ to record DA release in two striatal regions (Extended Data Fig. 1): the NAc core, vital to reward learning and addiction^18^; and the DLS, long linked to movement initiation and habit formation^19^. In one influential theory, demand for drugs of abuse begins with DA release in the NAc, which then spirals up to the DLS once behavior becomes habitual^20^. However, compared to the NAc, many fewer studies have recorded DA release in the DLS during reward tasks.

We aligned our recordings to the cue denoting reward availability, as this cue stayed identical in all conditions even as the cost and benefit varied. Thus, any observed changes in DA release could not be due to the sensory qualities of the stimulus, but rather to the perceived meaning of the associated reward. As expected, we found that increased sucrose concentration (Fig. 1d) and quantity (Fig. 1f) enhanced cued DA release in the NAc. DLS DA exhibited the same pattern (Fig. 1e and 1g), although the kinetics of the DA response in DLS was strikingly faster than that observed in the NAc.

Surprisingly, increased cost clearly enhanced DA responses in both regions (Fig. 1h, i), despite the cue and reward being held constant. This effect was not due to differences in the amount of time between rewards, because even when we kept this time interval constant, DA release increased with increasing FR (Extended Data Fig. 2a, b). To determine whether the effect of cost on DA release might relate to the order the FRs were presented, we conducted two additional experiments. In one, we ran mice through the five FRs on different days, and still found the same effect (Extended Data Fig. 2c, d). In a second control experiment, we ran new cohorts of mice through two separate sessions: one where FR1 preceded FR10, and another where FR10 preceded FR1. Regardless of order, DA release was larger for the higher FR (Extended Data Fig. 2e-h). We conclude that in both striatal regions, DA encodes both benefit and cost and that (sunk) cost positively correlates with DA release.

### Economic demand curves dissociate consumption from motivation

To make rational decisions, individuals must combine cost and benefit to determine if the benefit is worth the cost. This cost-benefit calculus is at the heart of motivation^21^. To measure motivation, we turned to the classic economic analysis of demand curves^22,23^. For each session, we plotted reward consumption as a function of cost (Fig. 2a), revealing two orthogonal parameters: Q_0_, the preferred level of consumption at no cost; and alpha, the sensitivity of consumption to cost (i.e., elasticity). Q_0_ is the y-intercept of the demand curve and is influenced predominantly by overall reward consumption or reward rate. In contrast, alpha, which depends on the demand curve’s slope, is inversely proportional to the subject’s motivation to obtain a reward^24^.

**Fig. 2:**
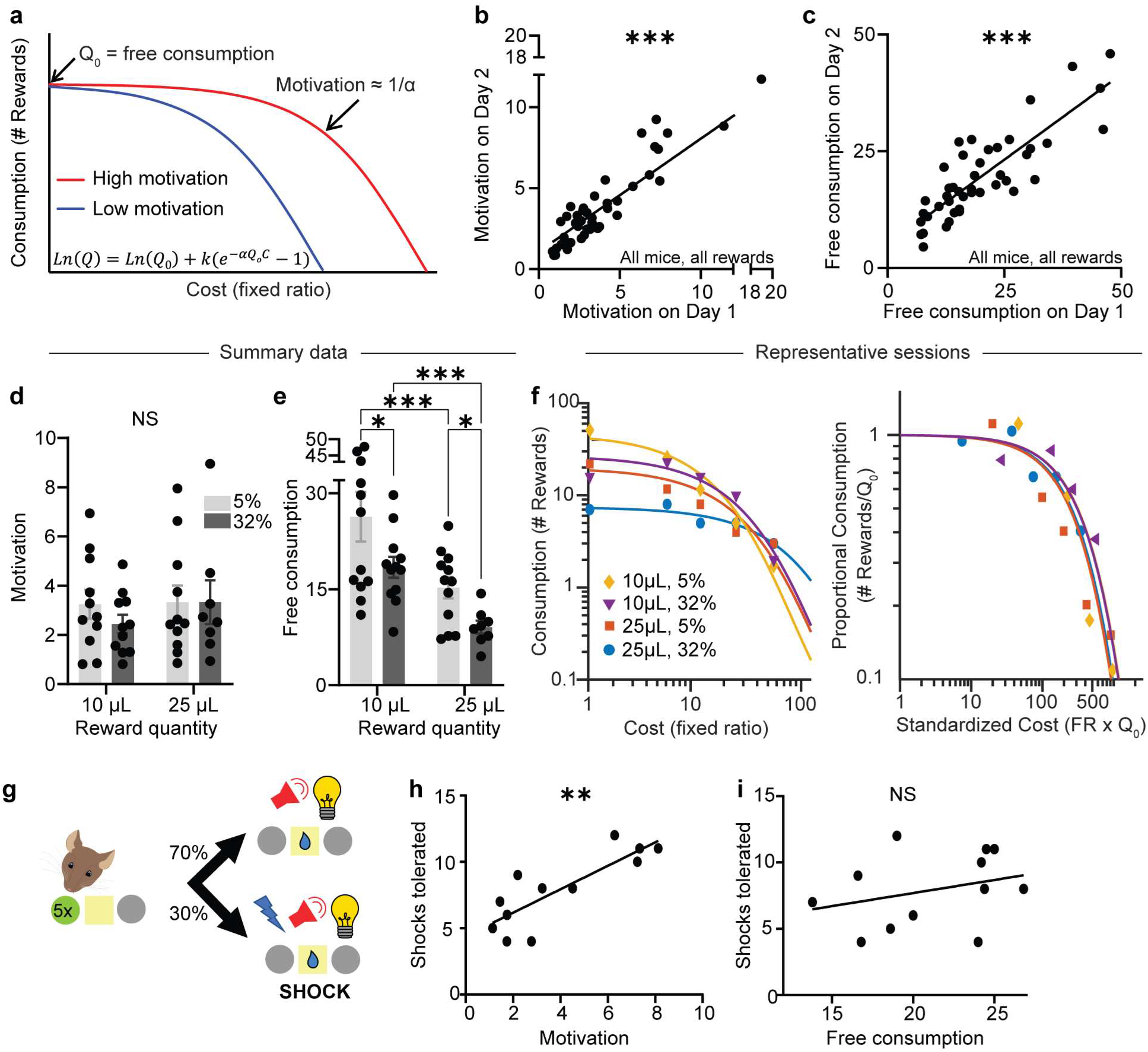
Economic demand curves dissociate consumption from motivation. **a**, Schematic demand curves plotting reward consumption as a function of cost. Shallower slope denotes higher motivation. **b**, Motivation parameter (1/α) on consecutive days with the same reward (Spearman correlation, r=0.86, P<0.0001; n=44 sessions from 12 mice). **c**, Free consumption parameter (Q0) on consecutive days with the same reward (Spearman correlation, r=0.82, P<0.0001; n=44 sessions from 12 mice). **d**, Average motivation as a function of sucrose quantity and concentration (Mixed-effects model, fixed effect of quantity, F_1,11_=3.95, P=0.073; fixed effect of concentration, F_1,11_=0.36, P=0.56; n=12). **e**, Average free consumption as a function of sucrose quantity and concentration (Mixed-effects model, fixed effect of quantity, F_1,11_=26.3, P=0.0003; fixed effect of concentration, F_1,11_=9.88, P=0.0093; Tukey-corrected multiple comparisons: 10μL-5% vs 10μL-32%, P=0.025; 10μL-5% vs 25μL-5%, P=0.0003; 10μL-32% vs 25μL-32%, P=0.0003; 25μL-5% vs 25μL-32%, P=0.025; n=12). **f**, Left, representative demand curves from a single mouse exposed to four rewards. Right, the same demand curves, now normalized by free consumption. **g**, Schematic of FR5 shock task. **h**, **i**, Previously-measured motivation (**h**: Spearman correlation, r=0.79, P=0.003; n=12) and free consumption (**i**: Spearman correlation, r=0.38, P=0.22; n=12) versus number of shocks tolerated. NS, not significant; *P<0.05, ***P<0.001. Error bars denote s.e.m.

To validate that our analyses accurately reflected the consistency in subjects’ behavior, we repeated the same demand curve assays with a constant sucrose reward on consecutive days and found that our metrics for motivation (Fig. 2b) and free consumption (Fig. 2c) remained consistent, with minor day-to-day variability. Furthermore, varying the reward magnitude did not cause changes in alpha, our measure of motivation (Fig. 2d), but did cause changes in Q_0_, the free consumption parameter (Fig. 2e). This dissociation between motivation and free consumption is expected^22^, and helps ensure that the alpha parameter does not simply reflect the dose of the reward, but rather the reward’s motivational value. In a typical subject, free consumption varied from session to session as different sucrose rewards were provided (Fig. 2f, left graph), but normalizing each demand curve by the free consumption (Fig. 2f, right graph) revealed that alpha, the slope of the curve and hence the motivational pull of sucrose, remained consistent despite changes in dose.

As a final test of the validity of the task, we asked whether the alpha parameter predicted another common metric for motivation: tolerance to punishment. In this new task, mice received sucrose reward at FR5 on 70% of trials, and in the remaining 30% of trials, they received both sucrose reward and a mild footshock (Fig. 2g). We found that the motivation parameter (1/alpha) calculated from previous demand curve assays (Fig. 2h), but not the free consumption parameter, Q_0_ (Fig. 2i), predicted the number of shocks a subject was willing to experience to obtain the reward. These results provide strong evidence that the behavioral economics task used in this study successfully measures motivation independently from dose or preferred consumption.

### Striatal DA release inversely reflects a reward’s motivational value

To determine how striatal DA release relates to motivation or reward consumption, we took advantage of the daily variation in subjects’ performance. Strikingly, DA release to a given fixed reward reflected motivation level (Fig. 3a, b) but not free consumption (Fig. 3c, d). In sessions when mice were more motivated for a given reward, less DA was released in both striatal regions (Fig. 3e, g and Extended Data Fig. 3). Furthermore, group comparisons revealed that mice with higher average motivation levels exhibited lower average DA responses to a fixed reward in both ventral and dorsal striatum (Fig. 3f, h). Thus, DA release reflects both day-to-day and individual-to-individual variability in motivation. In both cases, the lower the motivation, the higher the striatal DA release.

**Fig. 3:**
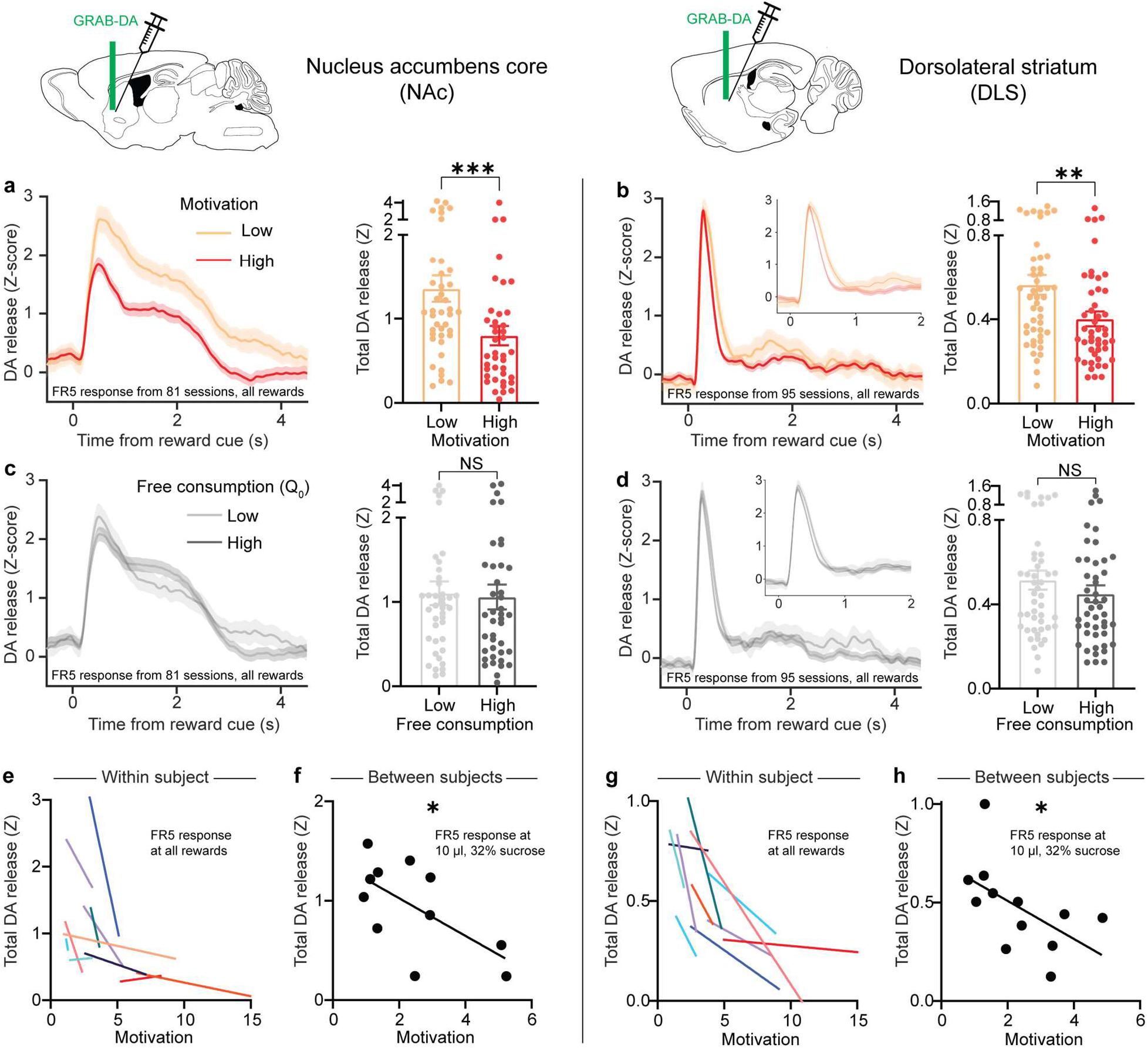
Striatal DA release inversely reflects a reward’s motivational value. **a-d**, Left, Z-scored photometry traces of GRAB-DA fluorescence in the NAc (**a**, **c**) or DLS (**b**, **d**) aligned to reward-predicting cues. Right, AUC of Z-scored photometry traces from 0 to 4s after cue onset as a function of motivation (**a**: Mann-Whitney test, U=475, P=0.001; n=81 sessions from 12 mice; **b**: Mann-Whitney test, U=735, P=0.0049; n=94 sessions from 12 mice) or free consumption (**c**: Mann-Whitney test, U=775, P=0.68; n=81 sessions from 12 mice, **d**: Mann-Whitney test, U=978, P=0.34, n=94 sessions from 12 mice). **e**, **g**, Motivation versus the AUC of cue-evoked GRAB-DA signal in the NAc (**e**) or DLS (**g**) for individual mice (each colored line denotes one mouse). **f**, **h**, Average motivation for 10 μL, 32% sucrose reward versus the average GRAB-DA signal for the cue predicting that reward in the NAc (**f**: Spearman correlation, r = −0.62, P=0.048; n=11) or DLS (**h**: Spearman correlation, r = −0.64, P=0.028; n=12). NS, not significant; *P<0.05, **P<0.01, ***P<0.001. Error bars denote s.e.m.

### Optogenetically-evoked striatal DA release is sensitive to cost

All of the above results rely on sucrose reward, which is sensitive to circuits associated with taste and satiety. To bypass these circuits and examine the pure motivational effect of striatal DA release, in a separate group of mice we employed optogenetic self-stimulation. Each DAT-Cre mouse received two viral injections: one to deliver either the excitatory opsin ChRMINE or the inert fluorophore mScarlet in DA neurons of the ventral tegmental area (VTA) or substantia nigra pars compacta (SNc), and another to deliver GRAB-DA in the appropriate target in the NAc or DLS. We then implanted a fiber over the NAc or DLS, allowing us to simultaneously stimulate DA inputs and record the resulting DA release. The task was identical as before, except that instead of poking for sucrose, mice poked for optogenetic stimulation of striatal DA release, accompanied by the same light-sound cue (Fig. 4a). As expected, mice learned to poke for DA input stimulation in both the NAc (Fig. 4b) and DLS (Fig. 4d), increasing both the number of rewards (left) and poking accuracy (right) over training. Control mice without opsin did not learn the task for light stimulation in either NAc (Fig. 4c) or DLS (Fig. 4e).

**Fig. 4:**
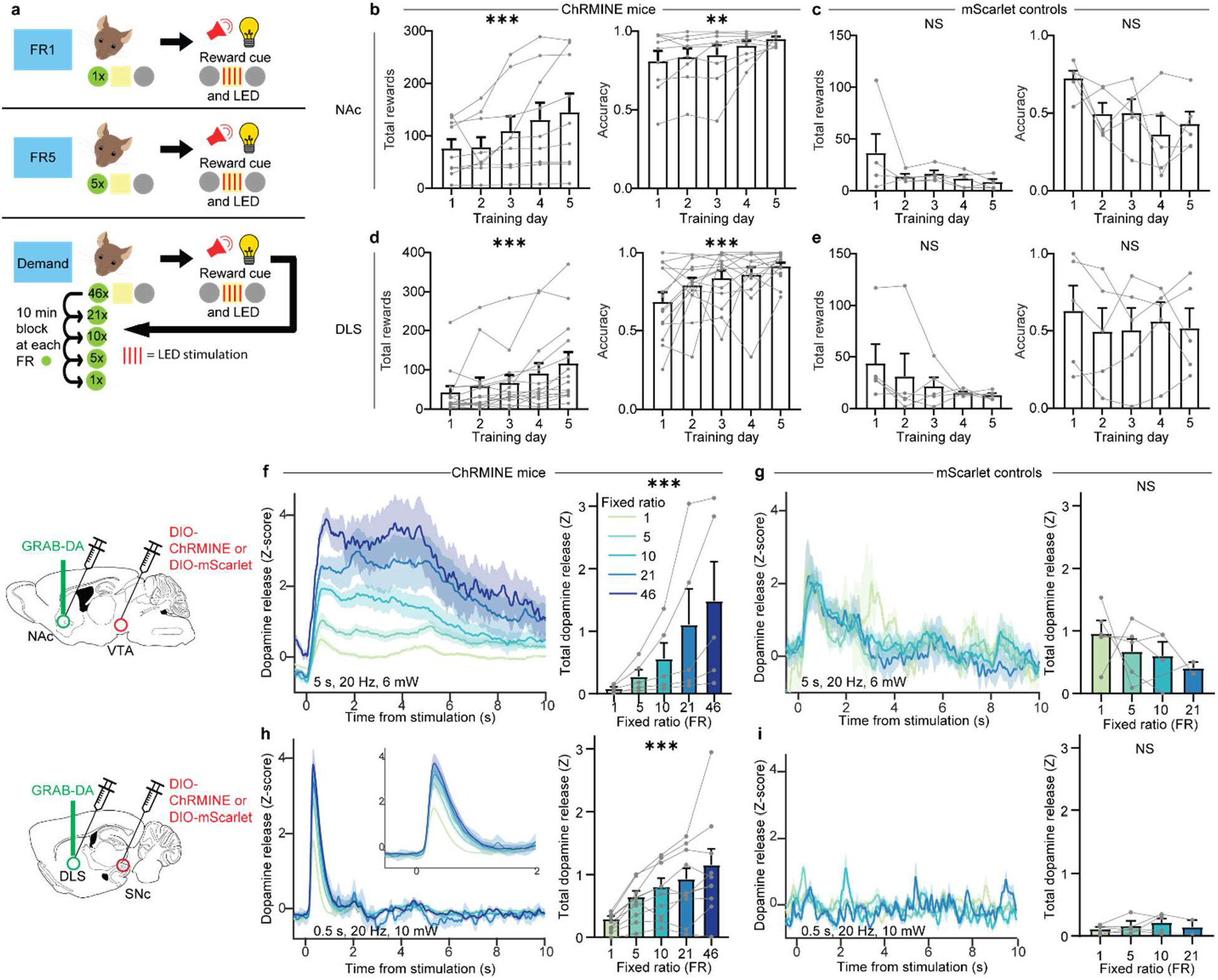
Optogenetically-evoked striatal DA release is sensitive to sunk cost. **a**, Schematic of optogenetic self-stimulation task. **b-e**, Number of rewards earned each day (left) and fraction of nosepokes in the active poke (right) during FR1 training in mice with ChRMINE in the NAc (**b**: rewards, Friedman test, Friedman statistic=31.9, P<0.0001; n=9; accuracy, Friedman test, Friedman statistic=15.3, P=0.0042; n=9) or DLS (**d**: rewards, Friedman test, Friedman statistic=31.2, P<0.0001; n=14; accuracy, Friedman test, Friedman statistic=19.8, P=0.0006; n=14), or control mice with mScarlet in the NAc (**c**: rewards, Friedman test, Friedman statistic=9.01, P=0.061; n=5; accuracy, Friedman test, Friedman statistic=9.12, P=0.058; n=5) or DLS (**e**: rewards, Friedman test, Friedman statistic=2.87, P=0.58; n=5; accuracy, Friedman test, Friedman statistic=1.92, P=0.75; n=5). **f-i**, Left, Z-scored photometry traces of GRAB-DA fluorescence in the NAc (**f**, **g**) or DLS (**h**, **i**) aligned to optogenetic stimulation in ChRMINE mice (**f**, **h**) or mScarlet controls (**g**, **i**). Right, AUC of Z-scored photometry traces from 0 to 10s after stimulation onset as a function of FR in mice with ChRMINE in the NAc (**f**: Friedman test, Friedman statistic=20.0, P=0.0005; n=5) or DLS (**h**: Friedman test, Friedman statistic=32.4, P<0.0001; n=10) or control mice with mScarlet in the NAc (**g**: Mixed-effects model, F_3,12_=0.44, P=0.44; n=5) or DLS (**i**: Mixed-effects model, F_3,8_=0.81, P=0.52; n=5). NS, not significant; *P<0.05, **P<0.01, ***P<0.001. Error bars denote s.e.m.

Despite using identical light stimulation parameters, as cost increased (i.e. the number of nose pokes required to receive stimulation), optogenetically-evoked DA release in both the NAc (Fig. 4f) and the DLS (Fig. 4h) increased. Control mice, who occasionally nose-poked despite receiving no reward, exhibited no relationship between cost and DA release (Fig. 4g, i). Although these results are consistent with the sucrose experiment (Fig. 1h, i), they were unexpected in that they suggest that the effort put forth by a subject dramatically influences the magnitude of optogenetically-evoked striatal DA release. Thus, we considered an alternative plausible interpretation: that the effect of varying the FR on optogenetic striatal DA release might reflect the longer interval between light stimulation during higher FRs. Perhaps at shorter time intervals, optogenetic stimulation of DA inputs depletes DA and reduces the amount of DA release for subsequent stimulations. However, when we analyzed trials with similar reward intervals at different FRs, the effect held (Extended Data Fig. 4a, b).

As a further test of the potential influence of the interval between optogenetic stimulations, we conducted an experiment under anesthesia, in which light stimulations were delivered at different intervals based on the average intervals for each FR in our task. Again, DA release was not influenced by the interval between light stimulations (Extended Data Fig. 4c, d). Finally, we considered if the order of FRs could explain the results; for example due to photobleaching of the sensor over the course of each session. We thus examined how DA release changed over time in a session where optogenetic stimulation was delivered without cost (Extended Data Fig. 4e, f) or with cost held constant (Extended Data Fig. 4g, h). In both cases, we found that DA release did not differ based on whether optogenetic stimulation was delivered at the beginning or the end of the session. Therefore, we conclude that DA release evoked by local optogenetic stimulation of DA inputs in both the NAc and DLS is greatly influenced by the effort put forth to gain that reward, i.e. the sunk cost.

### Optogenetically-evoked DA release is inversely related to motivation

Analysis of behavioral economic demand curves provides a platform for determining the intrinsic motivational value of striatal DA release. Similar to our experiments using sucrose as the reward, we took advantage of variations in motivation from day to day and mouse to mouse and compared optogenetically-induced DA release in high-versus low-motivation sessions. Consistent with previous results, DA release was greater during low motivation sessions in both the NAc and DLS (Fig. 5a, b and Extended Data Fig. 5). In contrast, the magnitude of free consumption did not affect optogenetic striatal DA release (Fig. 5c, d). The inverse relationship between motivation and optogenetic striatal DA release held both within each subject (Fig 5e, g) and between subjects (Fig. 5f, h). Thus, although DA release was required for animals to learn and perform the task (Fig. 4c, e), greater optogenetic striatal DA release reflected lower motivation.

**Fig. 5:**
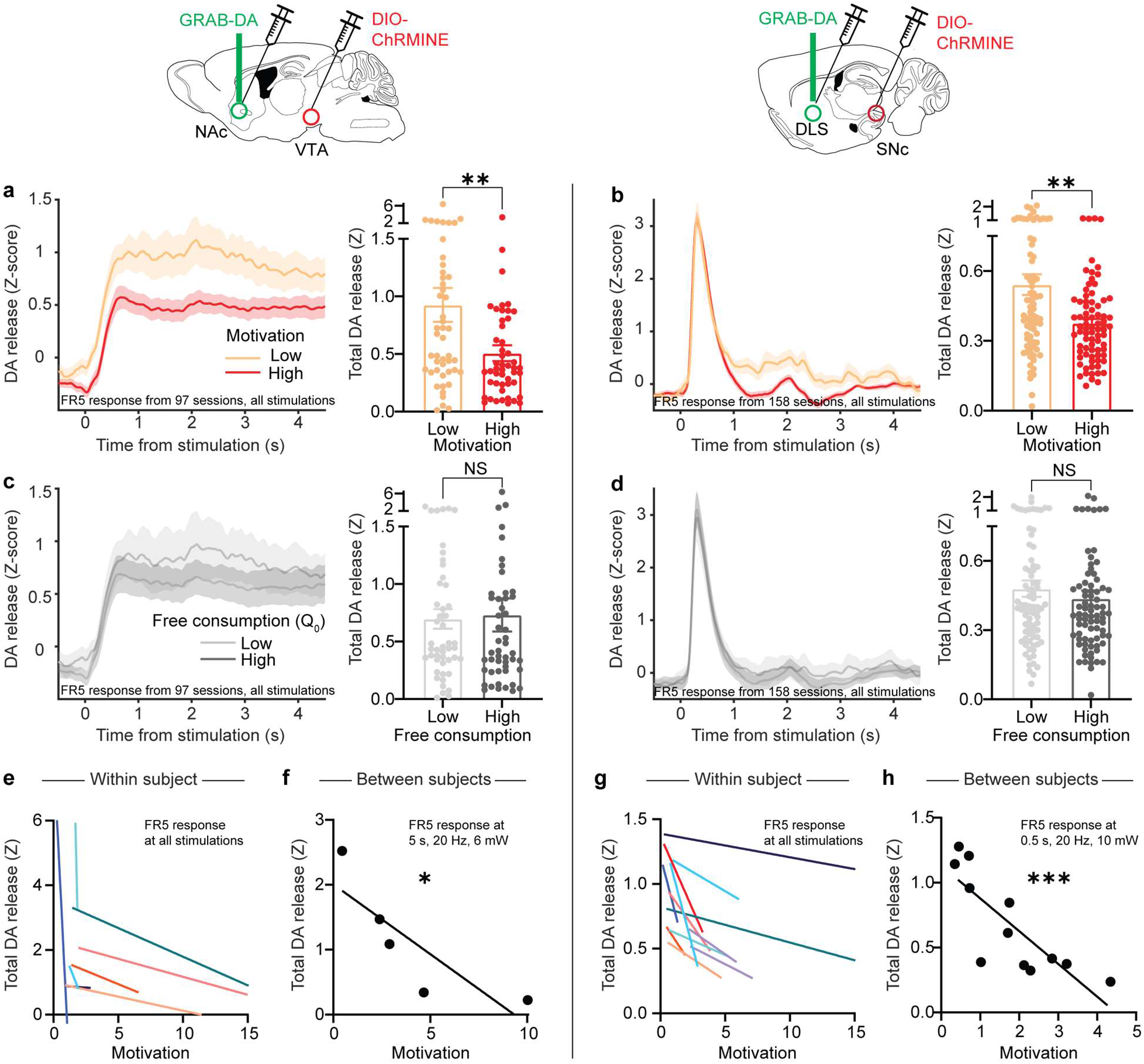
Optogenetically-evoked striatal DA release is inversely related to motivation. **a-d**, Left, Z-scored photometry traces of GRAB-DA fluorescence in the NAc (**a**, **c**) or DLS (**b**, **d**) aligned to optogenetic stimulation. Right, AUC of Z-scored photometry traces from 0 to 4s after stimulation onset as a function of motivation (**a**: Mann-Whitney test, U=798, P=0.0061; n=97 sessions from 9 mice; **b**: Mann-Whitney test, U=2293, P=0.0038; n=158 sessions from 14 mice) or free consumption (**c**: Mann-Whitney test, U=1085, P=0.52; n=97 sessions from 9 mice; **d**: Mann-Whitney test, U=2812, P=0.29; n=158 sessions from 14 mice). **e**, **g**, Motivation versus the AUC of stimulation-evoked GRAB-DA signal in the NAc (**e**) or DLS (**g**) for individual mice (each colored line denotes one mouse). **f**, **h**, Average motivation for optogenetic stimulation reward versus the average GRAB-DA signal for that reward in the NAc (**f**: Spearman correlation, r = −1.0, P=0.017; n=5) or DLS (**h**: Spearman correlation, r = −0.86, P=0.0006; n=12). NS, not significant; *P<0.05, **P<0.01, ***P<0.001. Error bars denote s.e.m.

### Testing the causal relationship between motivation and striatal DA release

By examining the behavioral variability that routinely occurs within and between subjects during our operant tasks, we provide evidence that the motivation to obtain a natural reward, sucrose, as well as the artificial “reward” of optogenetically-triggered striatal DA release, is inversely correlated with the magnitude of striatal DA release evoked by that reward. Our evidence thus far, however, cannot rule out an indirect or coincidental relationship between motivation and striatal DA release. To test the causal nature of this inverse relationship, we performed two further experiments. First, we sought to manipulate the intrinsic motivation for reward and observe the effect on striatal DA release elicited by that reward. To do so, we trained a new cohort of mice to nosepoke for sucrose reward and compared striatal DA release under two conditions: the standard condition, in which mice lack access to sucrose except for during the behavioral session; and a condition in which the animals had access to sucrose for 30 minutes before each behavioral session. As expected, prefeeding reduced motivation for sucrose (Fig. 6a, c) and these sessions were associated with higher striatal DA release in response to the same sucrose reward that was provided during standard sessions (Fig. 6b, d). Thus, a simple “natural” manipulation that reduced motivation for food without impairing task performance resulted in enhanced cued DA release in both the NAc and DLS.

**Fig. 6:**
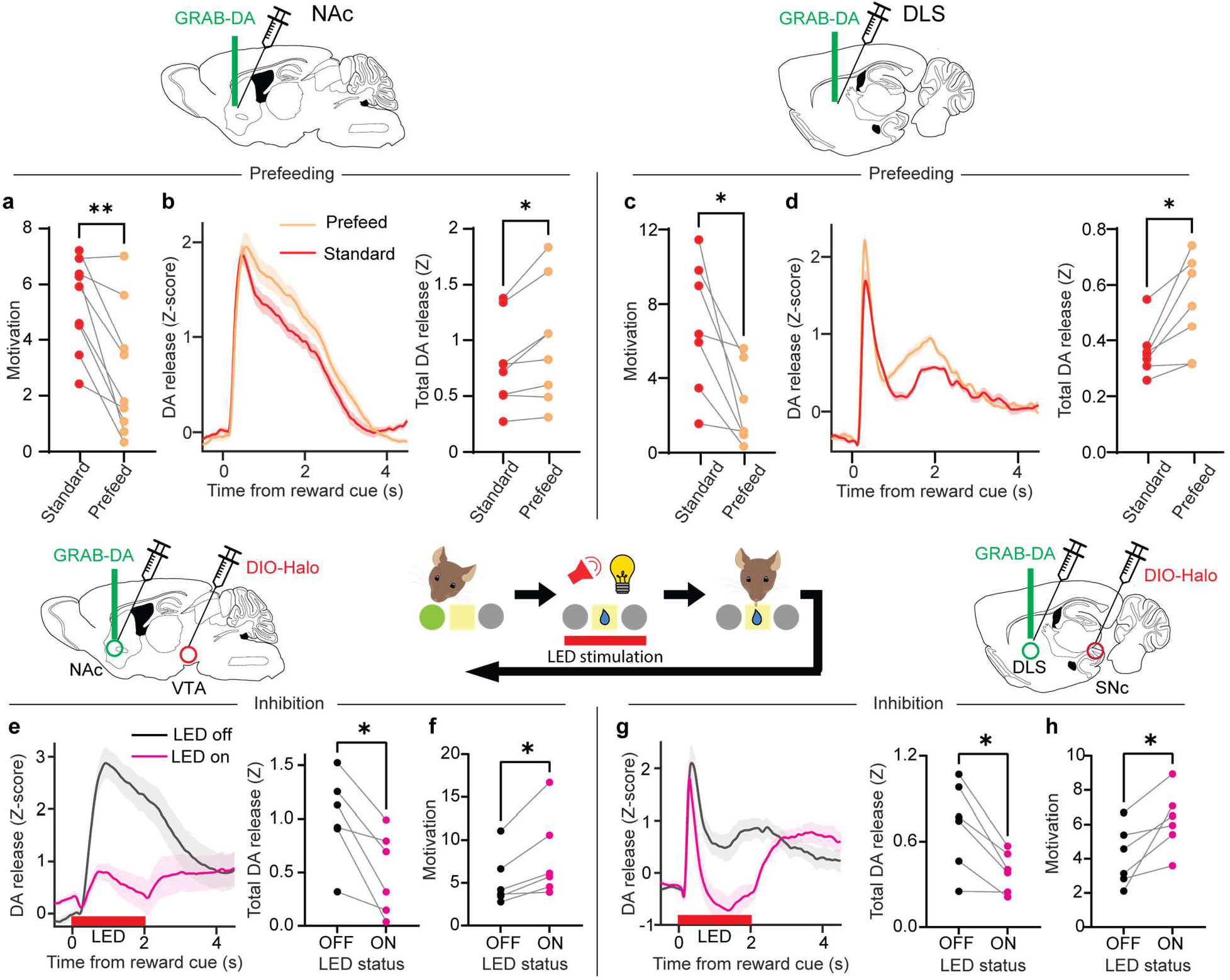
Bidirectional relationship between motivation and striatal DA release. **a**, **c**, Motivation for the same sucrose reward after 24-hr sucrose deprivation versus 30-min unlimited sucrose consumption for mice with fiber implants in the NAc (**a**: Wilcoxon matched-pairs signed rank test, W=-43, P=0.0078; n=9) or DLS (**c**: Wilcoxon matched-pairs signed rank test, W=-28, P=0.016; n=7). **b**, **d**, Left, Z-scored photometry traces of GRAB-DA fluorescence in the NAc (**b**) or DLS (**d**) aligned to reward-predicting cues. Right, AUC of Z-scored photometry traces from 0 to 4s after cue onset in standard versus pre-fed conditions in the NAc (**b**: Wilcoxon matched-pairs signed rank test, W=34, P=0.016; n=8) or DLS (**d**: Wilcoxon matched-pairs signed rank test, W=28, P=0.016; n=7). **e**, **g**, Left, Z-scored photometry traces of GRAB-DA fluorescence in the NAc (**e**) or DLS (**g**) aligned to cue onset in the optogenetic inhibition experiment. Right, AUC of Z-scored photometry traces from 0 to 4 s after cue onset as a function of whether inhibitory optogenetic stimulation was delivered during the cue in the NAc (**e**: Wilcoxon matched-pairs signed rank test, W=-21, P=0.031; n=6) or DLS (**g**: Wilcoxon matched-pairs signed rank test, W=-21, P=0.031; n=6). **f**, **h**, Motivation in sessions with or without optogenetic inhibition in the NAc (**f**: Wilcoxon matched-pairs signed rank test, W=21, P=0.031; n=6) or DLS (**h**: Wilcoxon matched-pairs signed rank test, W=21, P=0.031; n=6). *P<0.05, **P<0.01.

Second, we sought to manipulate the DA release elicited by a natural reward and observe the effect on motivation. A cohort of DAT-Cre mice received two viral injections: one to deliver the inhibitory opsin halorhodopsin (NpHR) in DA neurons of the VTA or SNc, and another to deliver GRAB-DA in the appropriate target in the NAc or DLS. We then implanted a fiber over the NAc or DLS, allowing us to simultaneously inhibit DA inputs and record the resulting DA release. Once they recovered from surgery, we trained these mice to nosepoke for sucrose reward in the same paradigm as before. After stabilizing their performance on the behavioral economics task, we used light to inhibit DA release during the 2-sec reward cue on each trial. We found that NpHR stimulation successfully reduced DA release in the NAc (Fig. 6e) and DLS (Fig. 6g), and that this manipulation increased motivation for sucrose reward (Fig. 6f, h). We conclude that there is a direct, bidirectional relationship between striatal DA release and motivation for natural reward.

## Discussion

Striatal DA release has long been linked to motivation, but most studies of DA and motivation have not recorded DA activity dynamics, and most DA recording studies have not measured motivation. Here we employed concepts from behavioral economics to gauge motivation for both a natural reward, sucrose, and the artificial reward of optogenetically-evoked striatal DA release, all while recording DA transients in the NAc and DLS, two striatal regions implicated in reward processing and operant behavior.

Our experiments generated several surprising results that help resolve controversies about the functional roles of striatal DA. First, striatal DA encodes <u>both</u> reward magnitude and sunk cost, which is the cost that has already been paid. Most prior studies have examined DA responses before the cost is paid, finding minimal cost sensitivity^9–14^ even though DA manipulations do appear to affect cost calculations^5–8^. In addition, influential recent work has debated whether DA activity dynamics better reflect reward value^25^ or changes in reward value^26^, but these studies did not modulate cost and thus did not address how DA release signals cost-benefit trade-offs. By varying cost and measuring DA release <u>after</u> the cost is paid, we demonstrate robust cost sensitivity, helping resolve the longstanding debate about whether striatal DA release is primarily important for reward benefit or cost. Our results, consistent with some recent work^27^, demonstrate that DA release encodes both. These results also provide a neural mechanism for the longstanding psychological observation that we value rewards more if we worked harder for them^28^.

Second, by leveraging individual and temporal variation in motivation, we demonstrate that there is a surprising inverse relationship between transient DA release and motivation. The higher the subject’s motivation, the lower the striatal DA release for a fixed reward, and vice versa. This finding may appear to contradict the commonly held notion that DA facilitates motivation^4,29^. However, it is consistent with the longstanding observation in stimulant addiction that the most severe drug users, who have high motivation to seek and ingest drugs of abuse, exhibit decreases in drug-elicited DA release, which predict the magnitude of drug use and craving^30,31^. Our findings suggest that chronic drug use, rather than simply triggering a dysphoric, low-DA withdrawal state^32,33^, may tap into a previously unknown mechanism that modulates striatal DA release depending on motivation. Furthermore, this mechanism appears to work in a qualitatively similar way in both the NAc and the DLS, although the two sites had different release kinetics, perhaps because of greater dopamine transporter expression^34^ and thus faster uptake^35^ in dorsal compared to ventral striatum.

A third surprising finding was that striatal DA release elicited by optogenetic stimulation of DA inputs was highly context dependent in that it too was influenced by costs, benefits and motivation. Since we only activated DA axons, these results support the importance of local modulation of DA release^36–39^ in addition to input integration at the level of DA cell bodies. They also emphasize that behavioral context can influence the neural consequences of “simple” optogenetic manipulations.

Our results support the proposal that striatal DA release encodes costs, benefits and motivation simultaneously but over different time scales. On a trial-by-trial basis, striatal DA release integrates benefit size and sunk cost. Motivation for rewards, in contrast, reflects a state that changes over slower time scales, consistent with recent work on hunger^40,41^, thirst^41,42^, and other internal homeostatic drives^43,44^. The striatal DA signals encoding reward are dampened when motivation is high and augmented when motivation is low. These results are consistent with an accelerator/brake model of DA action in which a high motivational state “releases the brake” such that a lighter push on the accelerator (i.e. smaller DA release) is necessary to initiate actions^45,46^. Future work that measures the downstream neural consequences of DA release in the striatum and other target regions may reveal how DA not only reflects motivation but also controls it.

## METHODS

### Mice

Female and male mice, first-generation offspring of female CD-1 mice (Charles River, strain 22) and either male C57BL/6J mice (Jackson Laboratory, strain #664) or male DAT:IRES-Cre mice on a C57BL/6J background (Jackson Laboratory, strain #006660) were used as experimental subjects. This CD-1/C57 hybrid strategy was chosen to facilitate related research on aggressive behavior as previously validated^47,48^. All transgenic animals used in these experiments were genotyped and found to be heterozygous for Cre recombinase. Mice were group-housed before surgical procedures. After surgery, males were single-housed to reduce aggression within a cage. Behavioral experiments were conducted during the dark cycle (lights off at 09:00, on at 21:00) when mice were 10-24 weeks old. Mice were maintained under food restriction for all behavioral experiments (>80% body weight from baseline ad libitum period). All procedures complied with the animal care standards set forth by the National Institutes of Health (NIH) and were approved by the Stanford University Administrative Panel on Laboratory Animal Care and Administrative Panel of Biosafety.

### Stereotactic injections and viruses

Mice (8-12 weeks of age) were anesthetized with isoflurane (5% induction, 1-2% maintenance). Meloxicam (5 mg/kg) was administered subcutaneously at the start of surgery and 24 and 48 hours after. Heads were fixed on a Kopf stereotaxic apparatus, a small incision was made in the scalp, and burr holes were drilled in the skull over the sites of injection. The following bregma coordinates were used: VTA, −3.1 mm anteroposterior (AP), ±0.7 mm mediolateral (ML), 4.2 mm dorsoventral (DV) from dura; SNc, −3.1 mm AP, ±1.0 mm ML, 4.3 mm DV; NAc, 1.5 mm AP, ±0.9 mm ML, 4.2 mm DV; DLS, 0.8 mm AP, ±2.3 ML, 3.1 mm DV. Microliter syringes (Hamilton) were lowered to the specified depth from the dura and used to inject 0.5 μL of virus solution at a flow rate of 0.1-0.25 μL min^-1^. For experiments with two viral injections in the same location (e.g., for optogenetic stimulation and photometry recording), a total of 1 μL mixed solution was injected. Borosilicate optic fibers for photometry and/or optogenetic stimulation with 400-μm diameter and numerical aperture 0.66 (Doric) were implanted directly above the striatal virus injection site, ipsilateral to the midbrain virus injection site, and secured to the skull using light-cured dental adhesive cement (Geristore A&B paste, DenMat). For the cohort with recordings in both NAc and DLS (Extended Data Fig. 1a), the NAc and DLS fibers were implanted in different hemispheres. For other cohorts, the hemisphere was counterbalanced between mice.

Adeno-associated viruses (AAVs) used for stereotactic injections were AAV9-hSyn-DA2m (DA4.4, WZ Biosciences), AAV-8-EF1α-DIO-ChRmine-mScarlet-WPRE (Stanford Gene Vector and Virus Core), AAV-8-EF1α-DIO-mScarlet-WPRE (Stanford Gene Vector and Virus Core), and AAV-DJ-EF1α-DIO-NpHR3.0-mCherry (Stanford Gene Vector and Virus Core). AAV titres ranged from 1 × 10^12^ to 2 × 10^13^ gc ml^-1^.

### Optogenetic manipulations

For photostimulation of ChRMINE or NpHR, optical implants were connected to a 625nm LED light source (Prizmatix) via a plastic fiber (1mm diameter, NA 0.63) and a fiber optic rotary joint (Doric). In an attempt to mimic DA release dynamics for natural reward, we used slightly different sets of ChRMINE stimulation parameters for NAc and DLS. For the data presented in Fig. 4 and Extended Data Fig. 4, NAc stimulation was kept at 5 s, 20 Hz and 6mW, while DLS stimulation was kept at 0.5 s, 20 Hz and 10 mW, all with a pulse width of 10 ms. On other demand sessions (data shown in Fig. 5 and Extended Data Fig. 5), frequency was varied from 1-20 Hz, duration from 0.5-30 seconds and power from 3-10 mW, while pulse width was kept at 10ms. We found that these parameter changes did not systematically affect either motivation or DA release in the demand task, and thus we combined all sessions for the analyses in Fig. 5a-d, e, g and Extended Data Fig. 5. For NpHR stimulation, light was turned on (10 mW at fiber tip) for 2 s, only during the cue period.

### Fiber photometry

For photometry recordings, optical implants were connected to low-autofluorescence patch cords (400 μm diameter, 0.57 numerical aperture, Doric) via a ceramic sleeve (Doric). Signals then passed through a fiber optic rotary joint (Doric) before filtering through a fluorescence mini cube (Doric) and ultimately reaching a femtowatt photodetector (2151; Newport). Frequency-modulated blue (465 nm) and UV (405 nm) LEDs (Doric) were used to stimulate GRAB-DA emission and control signals through the same fibers. Blue LED power was adjusted to ~30μW at the fiber tip to ensure that the photodetector was not being saturated. Digital signals were sampled at 1.0173 kHz, demodulated, lock-in amplified, and recorded by a real-time signal processor (RZ5P, Tucker-Davis Technologies) using Synapse software (Tucker-Davis Technologies). Synchronized behavioral events from the Med Associates boxes were collected directly into the RZ5P digital input ports. To reduce any confounds from photobleaching, animals were recorded for about 5 min before behavioral testing began.

### Behavioral training

Behavioral training began at least 2 weeks after virus injections. For the first 2 days, mice were tethered to the fiber optic patch cord and allowed to explore the behavioral chambers (Med Associates) for 30 minutes. Each chamber had a speaker for white noise and three identical nose ports, each with an associated light and an infrared emitter-sensor to measure port entry times. The central nose port was defined as the reward port and was equipped to receive 10-25 μL sucrose (at 5% or 32% concentration) from a syringe pump.

After habituation to the chamber and tether, mice were exposed to 2 days of a Pavlovian task, in which rewards were delivered at random intervals between 25 and 35 s for 30 min. Each reward was delivered in tandem with a 2-s light-sound cue (white noise and central port light), with the goal that the mice would begin to associate the cue with reward delivery. Mice were then trained on FR1, meaning that a single poke in the active port was required for reward delivery. The identity of the active port (left or right) was kept consistent for each mouse, but counterbalanced between mice. For at least the first 3 days, or until the mice achieved 10 rewards per session, the active port was baited with a fruit loop to encourage the mice to explore that port. The bait was then removed and mice continued with at least five days of FR1, or until the number of rewards per session remained within 20% for 3 consecutive days. The training data in Fig. 1 derives from these unbaited FR1 sessions. Mice were then trained on FR5 until they achieved at least 10 rewards per session for 3 consecutive days, and then moved onto demand sessions (see below).

Training for optogenetic self-stimulation (Figs. 4-5) was identical, except instead of sucrose reward being delivered at the time of the light-sound cue, mice received optogenetic stimulation. Recording data from the optogenetic Pavlovian and FR5 sessions are shown in Extended Data Fig. 4e-g.

Male and female mice were used in approximately equal numbers in all behavioral tasks and there was no sex difference in any experimental condition. Subject mice were excluded from the analysis (less than 5% of 86 total) if they did not reach behavioral criteria or if on the basis of histological analysis, they had: (1) off-target transgene expression; (2) weak transgene expression; or (3) inaccurate implant placement.

### Demand sessions

During demand sessions, mice experienced 5 consecutive 10-min blocks, each with a different FR, in the following order: FR46, FR21, FR10, FR5, FR1. This order was chosen after extensive piloting showed that random presentation of FRs produced erratic responses, which is consistent with prior work^49^. We chose not to use an ascending order of FRs to reduce the chance of satiety from rapid consumption of rewards early in a session. In control experiments, we confirmed that the demand curves extracted from single sessions were similar to those extracted over 5 days, with a different FR presented on each day (data not shown).

On each trial, mice poked the required number of times in the active port and on the last required poke, the 2-s light-sound cue was presented. In the sucrose task, cue onset was simultaneous with sucrose delivery into the central port. After the cue ended and mice entered the central port to consume the sucrose, a light turned on in the active port denoting that another trial had begun and mice were free to poke again. In the optogenetic task, cue onset was simultaneous with LED stimulation. As soon as both LED stimulation and the cue had ended, a light turned on in the active port denoting that another trial had begun and mice were free to poke again.

We continued demand sessions for a single reward type for at least 2 days, or until the alpha parameter achieved stability within 20%. We then switched the reward type (i.e., sucrose volume or concentration or LED power, frequency or duration). The order of reward types was counterbalanced between mice.

### Shock experiment

After mice had completed the sucrose demand sessions, they returned to the standard FR5 task for one day, and on the next day performed the FR5-shock task. In this task, mice received reward as usual after 5 pokes, but on a randomly-selected 30% of trials, they also received a mild shock (0.2 mA, 0.5 s) through an electrified grid floor (Med Associates).

### Prefeeding experiment

Mice were trained to stability on the demand task as above, having access to sucrose only during the task itself. They then underwent a prefeeding procedure as follows: 30 min before the session, they were given access to 1.5 mL sucrose in their home cage. If they finished the sucrose, they were given another 0.5 mL, and if they finished that, they were given another 0.5 mL, and so on until the session was scheduled to begin. We found that this graded approach maintained the subjects’ motivation to perform the task better than providing unlimited sucrose (data not shown).

### Optogenetic inhibition experiment

DAT-Cre mice were injected with Cre-dependent NpHR3.0 in the VTA or SNc and GRAB-DA in the NAc or DLS, respectively, and a fiber was implanted above the striatal target. After 2 weeks of recovery, mice were trained on the sucrose demand sessions as usual, without any optogenetic stimulation, until the alpha parameter stabilized within 20% on 2 consecutive days. They then underwent inhibition experiments, which were identical except for the addition of 2-s red light stimulation during each cue period. Mice underwent at least 3 optogenetic sessions, until the alpha parameter stabilized within 20%. Data in Fig. 6e-h are derived from the last 2 days of standard performance and the last 2 days of optogenetic stimulation, when performance was stable.

### Alternating FR experiment

Mice were trained on the sucrose FR1 and FR5 tasks as usual and were then split into two cohorts. One cohort performed a task with 4, 10-min blocks in the following order: FR10, then FR1, then FR1, then FR10. The other cohort performed a task with the following order: FR1, then FR10, then FR10, then FR1. Each cohort repeated the task on 2 consecutive days and the data in Extended Data Fig. 2e-h comes from the average of the 2 days. We designed these tasks to compare cued DA responses on consecutive blocks as a function of FR order (1 before 10 or 10 before 1). Our design allows for both within-subject comparisons (analyzing the first 2 blocks vs the last 2 blocks of a session) and between-subject comparisons (analyzing the first 2 blocks in one cohort vs the other cohort).

### Anesthetized recording experiment

DAT-Cre mice were injected with Cre-dependent ChRMINE in the VTA or SNc and GRAB-DA in the NAc or DLS, respectively, and a fiber was implanted above the striatal target. After 2 weeks of recovery, mice were anesthetized with ketamine (60 mg/kg) and dexmedetomidine (0.6 mg/kg) through intraperitoneal injection, placed on a heating pad to maintain body temperature, and tethered to the photometry setup as usual. We then recorded GRAB-DA signals while mice received 60 optogenetic stimulations (for NAc: 5 s, 20 Hz, 6mW; for DLS: 0.5 s, 20 Hz, 10 mW) at intervals chosen at random from the following options: 5 s, 10 s, 20 s, 30 s, 70 s and 120 s. These intervals were chosen because they were the average intervals between rewards during the optogenetic demand task (NAc: 5s, 10s, 30s, 70s, 120s; DLS: 10s, 20s, 30s, 70s, 120s).

### Immunohistochemistry

Mice were perfused with 4% PFA and brains were removed and post-fixed overnight at 4 °C. 70–75-μm coronal sections were prepared on a vibratome and free-floating sections were processed for immunohistochemistry. After three 10-min washes in PBS on a shaker, the tissue was incubated with blocking solution (5% normal goat serum and 0.3% Triton X-100 in PBS) for 1 hour and then incubated in primary antibodies overnight at 4 °C on a shaker. The primary antibodies used were: rat mCherry monoclonal antibody 1:1,000 (Invitrogen, M11217) and chicken anti-GFP 1:1,000 (Aves labs, GFP-1020). After three washes of 10 min in PBS, secondary antibodies were added and incubated for 2 h at room temperature on a shaker. The secondary antibodies used were: goat anti-rat Alexa Fluor 594 1:750 (Invitrogen, A11007) and goat anti-chicken Alexa Fluor 488 1:750 (Abcam, ab150169). After three more washes, the slices were mounted with DAPI Fluoromount-G mounting medium (SouthernBiotech) onto microscopy slides for visualization at 4x using a Keyence BZ-X810 slide scanner (Keyence Plan Apo Chromat 4× objective, NA 0.20, filter cubes: DAPI, GFP, TRITC, wavelength: 430–741 nm, bit depth of images: 14 bit).

### Data analysis

Custom scripts in MATLAB (MathWorks) were used for signal processing. Signals were down-sampled 10x, underwent Loess smoothing (window size = 30 ms) to reduce high-frequency noise, and analyzed in 16-s windows around each cue delivery. Z-scores of the fluorescence for each trial were calculated based on the mean and standard deviation of the local baseline signal before each cue (−8 to −4 s before cue onset). Trials were combined into a single session average, and sessions were then averaged together for a single mouse. All photometry figures in the manuscript show the mean and standard error of the photometry signal across mice, a more conservative approach than using the trial- or session-average. To quantify total DA release, area under the curve was calculated using trapezoidal numerical integration on the Z-scores for the windows defined in the figure legends. To compare high-vs low-motivation sessions (Fig. 3a-d; Fig. 5a-d), we performed a median split by the alpha parameter.

To minimize bleaching confounds, we removed the first ~5 minutes of each recording, when the steepest bleaching was likely to occur. We also analyzed short windows of data with local baselines, avoiding any analysis of longer-timescale changes, which are more likely to be confounded by bleaching or other gradual changes (e.g., slight adjustments in the connection between the implant and the patch cord). In addition, we took the approach reported by Sych et al^50^ and examined simultaneously-recorded control signals (405 nm), which we found to be flat or slightly negative traces with substantially lower amplitude than what we observed with the experimental excitation (465 nm). Although this analysis is imperfect because 405 nm is not the isosbestic point for GRAB-DA^51^, the result implies that motion artifacts, bleaching, or other intrinsic, non-DA dependent signal changes could not have made a major contribution to our results.

Finally, we repeated all of our analyses using two methods to correct for bleaching, and our results held regardless of method. In the first method, the entire session was debleached according to the iterative method outlined by Bruno et al^52^, which calculates the ΔF/F in short moving windows, centers and normalizes these windows, and then repeats these calculations for 100 temporally-offset windows in the same session. The final debleached signal is the average of the 100 iterations. In the second method, we used polynomial curve fitting to fit the control 405 signal to the experimental 465 signal, then subtracted the fitted 405 signal from the experimental signal, and based all subsequent analyses on this subtracted signal. We chose not to complete this as our primary analysis because 405 nm is not the isosbestic point for GRAB-DA and because we observed that the bleaching rates for the two signals were often poorly aligned.

To produce demand curves, we used the nonlinear least squares method to fit behavioral data from each session to the following exponential equation^22^:

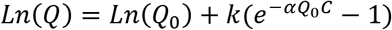

where Q is the number of rewards earned in a block, C is the cost (FR) of each reward for that block, and k is a constant that specifies the range of Q (here we set k = 3.2). From these fits we extracted estimates of 2 parameters: Q_0_, or the preferred consumption at no cost; and α, the slope or elasticity of the curve. We then converted α to our measure of motivation, also known as essential value, with the following equation^24^:

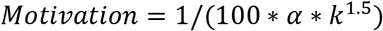

As in prior work^23^, goodness of fit was calculated with R^2^ and sessions were included in analysis only when R^2^ > 0.3. The median R^2^ ranged between 0.88 and 0.94 for the different behavioral cohorts.

### Statistical methods and reproducibility

All experiments were performed without knowledge of the virus that had been injected or the transgene being expressed, and were analyzed without knowledge of the specific manipulation each mouse had undergone. No statistical methods were used to predetermine sample size, which was based on previous experience with the variance of the assays. All data were tested for normality of sample distributions and when violated, non-parametric statistical tests were used. Friedman’s test was used to compare three or more matched groups (e.g., DA response from the same animals across 5 FRs). If individual data points were missing from these matched comparisons, mixed-effects models were used instead. Mixed-effects models were also used when examining the effects of multiple fixed effects (e.g., sucrose quantity and concentration) accounting for the random effect of subject. In these cases, the significance of the fixed effects was reported in the figure legends and if there were significant fixed effects, Tukey-corrected post-hoc comparisons were reported with asterisks in the figures. Wilcoxon matched-pairs signed rank tests were used to compare two paired groups (e.g., DA response from the same animals across two concentrations of sucrose). Spearman correlations were used to measure the association between two independently-measured observations (e.g., motivation on consecutive days). Mann-Whitney tests were used to compare two unmatched groups (e.g., DA response in high-motivation vs low-motivation sessions). Kruskal-Wallis tests were used to compare three or more unmatched groups (e.g., DA responses from trials with different inter-reward intervals). All statistical tests were two-sided and performed in MATLAB (Mathworks) or Prism (GraphPad). NS, not significant. *P<0.05, **P<0.01, ***P< 0.001. In all figures, data are shown as mean+ s.e.m.

## ACKNOWLEDGMENTS

We thank Marija Kamceva, Zihui Zhang, Anjali Temal, Ariana Rodrigues and members of the STAAR and Malenka labs for technical assistance and discussions. The Stanford Gene Vector and Virus Core provided reagents. This work was supported by philanthropic funds donated to the Nancy Pritzker Laboratory at Stanford University. N.E. was supported by NIH grant K08 MH123791, a Brain & Behavior Research Foundation Young Investigator Grant, a Burroughs Wellcome Fund Career Award for Medical Scientists, a Stanford Society of Physician Scholars Research Grant and the Simons Foundation Bridge to Independence Award. G.T. and A.R.W. were supported by Stanford MedScholars grants. D.C.P. was supported by an NSF Graduate Research Fellowship and an HHMI Gilliam Fellowship (with R.C.M.). R.C.M. was supported by a Stanford University Wu Tsai Neurosciences Institute grant and NIH grant P50 DA042012.

## AUTHOR CONTRIBUTIONS

N.E. and B.S.B. conceived the study with support from R.C.M. N.E. designed the experiments with input from R.C.M. N.E. and R.C.M. interpreted the results and wrote the paper, which was edited by all authors. A.R.W. designed and performed the sucrose photometry experiments. G.C.T. designed and performed the inhibition and prefeeding experiments. G.C.T., A.K.O., A.N.S., A.M.G., and L.T. performed the optogenetics experiments. D.C.P. and J.T. contributed to the design and analysis of the optogenetics experiments. D.C.P. performed the histological verifications of fiber placements.

## DATA AVAILABILITY

The datasets generated and analyzed during this study are included in this article and available from the corresponding authors upon reasonable request.

## CODE AVAILABILITY

Code used for data processing and analysis is available from the corresponding authors upon reasonable request.

## ADDITIONAL INFORMATION

Correspondence and requests for materials should be addressed to Neir Eshel

and Robert C. Malenka.

---

**Extended Data Fig. 1:**
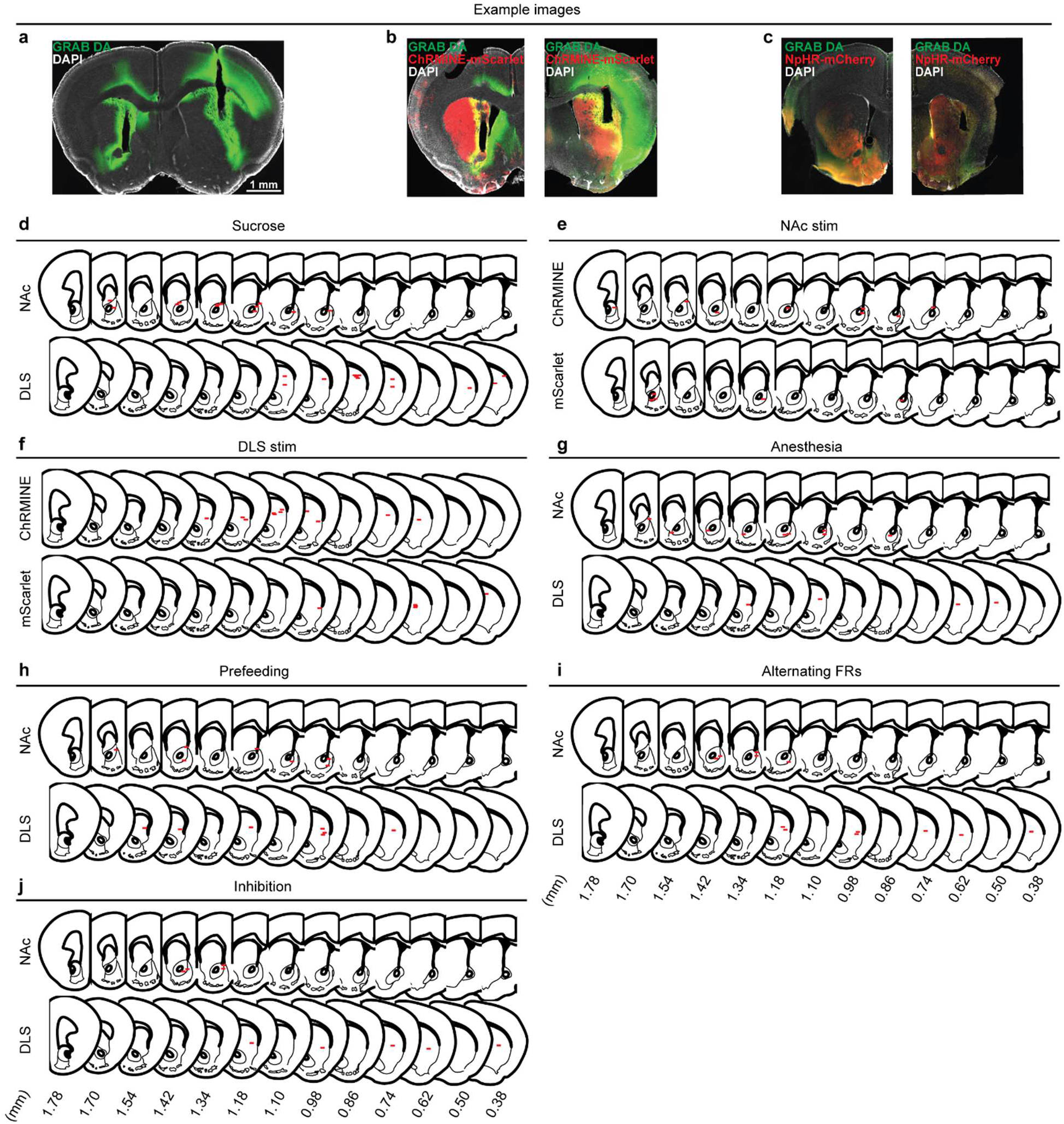
Histological validation of fiber implant targeting. **a**, Bilateral fiber implants and GRAB-DA expression in a representative mouse used in the sucrose experiments. **b**, Fiber implants and GRAB-DA and ChRMINE expression in the NAc (left) and DLS (right) for representative mice in the optogenetic stimulation experiments. **c**, Fiber implants and GRAB-DA and NpHR expression in the NAc (left) and DLS (right) for representative mice in the optogenetic inhibition experiments. **d**, Fiber implant tip locations in the NAc (top) and DLS (bottom) of mice used in the sucrose experiments. **e**, Fiber implant tip locations in the NAc of mice expressing ChRMINE (top) or mScarlet (bottom) used in the optogenetic stimulation experiments. **f**, Fiber implant tip locations in the DLS of mice expressing ChRMINE (top) or mScarlet (bottom) used in the optogenetic stimulation experiments. **g**, Fiber implant tip locations in the NAc (top) or DLS (bottom) of mice used in the anesthesia experiments. **h**, Fiber implant tip locations in the NAc (top) or DLS (bottom) of mice used in the prefeeding experiment. **i**, Fiber implant tip locations in the NAc (top) or DLS (bottom) of mice used in the alternating FRs experiments. **j**, Fiber implant tip locations in the NAc (top) or DLS (bottom) of mice used in the optogenetic inhibition experiments. Mice used in more than one experiment have their fiber placement repeated in each experiment.

**Extended Data Fig. 2:**
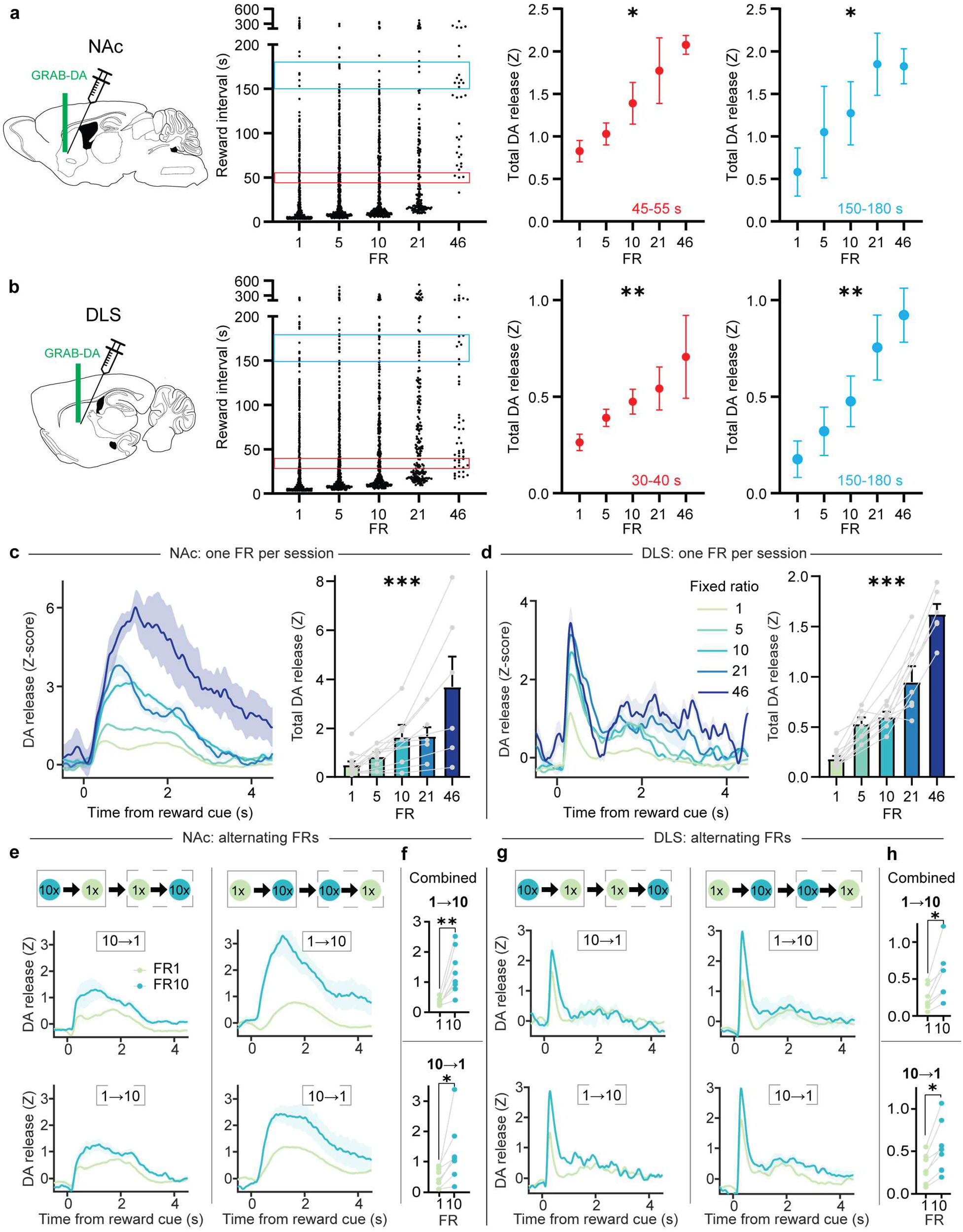
The effect of FR on DA release does not depend on inter-reward interval or the order of FRs. **a, b,** Left, Distribution of inter-reward intervals as a function of FR for mice with fibers in the NAc (**a**) or DLS (**b**). Middle, Right, AUC of GRAB-DA photometry traces in response to reward cues on trials with short (Middle, **a**: Kruskal-Wallis test, Kruskal-Wallis statistic=9.97, P=0.04; n=12; **b**: Kruskal-Wallis test, Kruskal-Wallis statistic=10.99, P=0.027; n=12) or long (Right, **a**: Kruskal-Wallis test, Kruskal-Wallis statistic=14.5, P=0.006; n=12; **b**: Kruskal-Wallis test, Kruskal-Wallis statistic=15.9, P=0.003; n=12) inter-reward intervals. **c**, **d**, Left, Z-scored photometry traces of GRAB-DA fluorescence in the NAc (**c**) or DLS (**d**) aligned to cue onset in sessions with a single FR. Right, AUC of Z-scored photometry traces from 0 to 4s after cue onset as a function of FR in the NAc (**c**: Mixed-effects model, F_4,20_=9.47, P=0.0002; n=12) or DLS (**d**: Mixed-effects model, F_4,20_=73.1, P<0.0001; n=12). **e**, **g**, Left, Z-scored photometry traces of GRAB-DA fluorescence in the NAc (**e**) or DLS (**g**) for mice performing a session with FRs presented in the following order: 10, then 1, then 1, then 10. Right, Z-scored photometry traces for a separate cohort of mice that performed a session with the following FR order: 1, then 10, then 10, then 1. Top, fluorescence traces for the first two FRs presented in a session. Bottom, fluorescence traces for the last two FRs presented in a session. **f**, **h**, Top, AUC of Z-scored photometry traces in the NAc (**f**: Wilcoxon matched-pairs signed rank test, W=36, P=0.0078; n=8) or DLS (**h**: Wilcoxon matched-pairs signed rank test, W=28, P=0.016; n=7) for consecutive blocks in which FR1 is presented before FR10. Bottom, AUC of Z-scored photometry traces in the NAc (**f**: Wilcoxon matched-pairs signed rank test, W=28, P=0.016; n=7) or DLS (**h**: Wilcoxon matched-pairs signed rank test, W=28, P=0.016; n=7) for consecutive blocks in which FR10 is presented before FR1. Regardless of FR order or timing within a session, FR10 trials always elicited greater DA release than FR1 trials. *P<0.05, **P<0.01, ***P<0.001.

**Extended Data Fig. 3:**
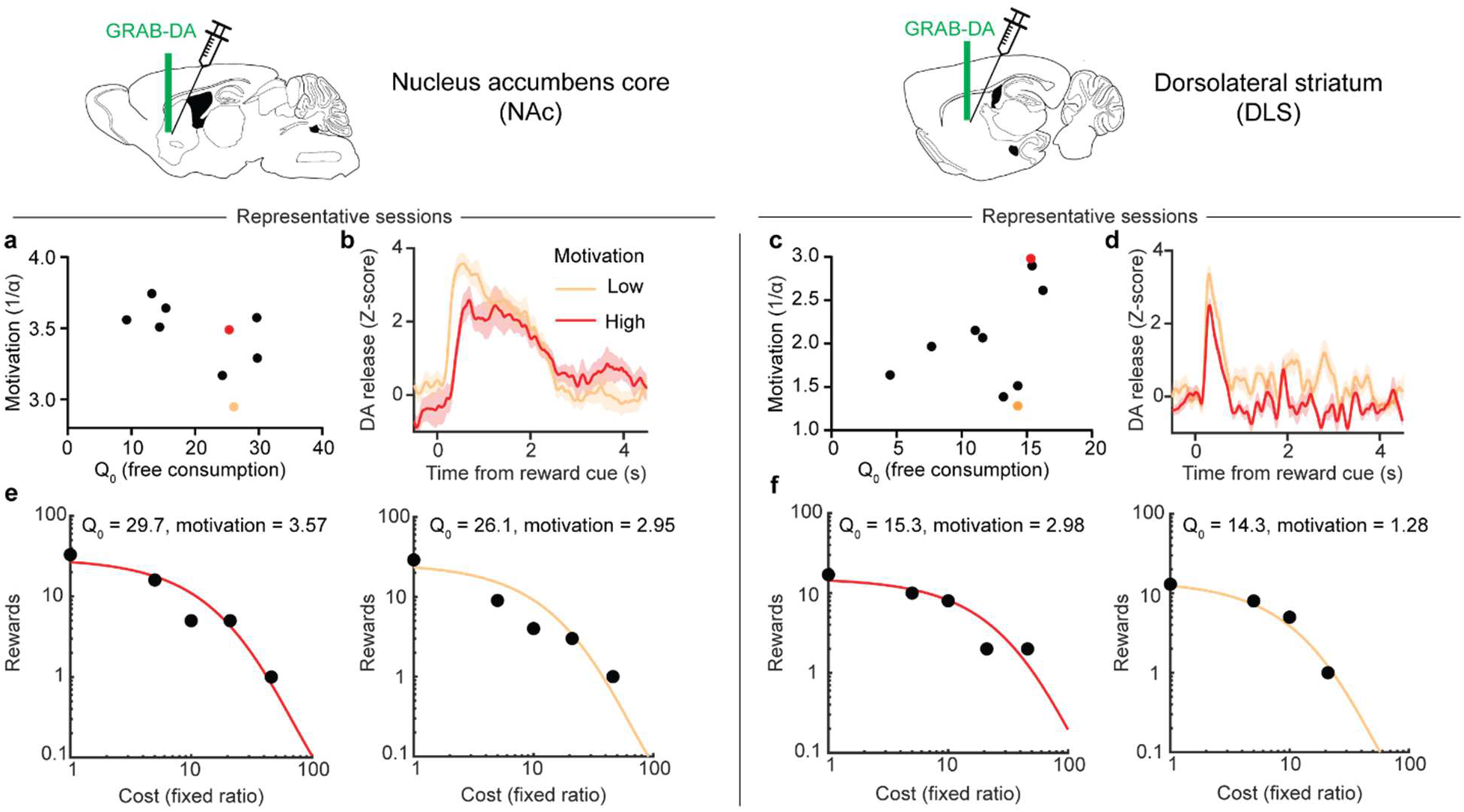
Representative examples of the inverse relationship between motivation and DA release for sucrose rewards. **a**, **c**, Free consumption parameter versus motivation parameter extracted from individual sessions in one representative mouse with recordings in the NAc (**a**) or DLS (**c**). **b**, **d**, Z-scored photometry traces of GRAB-DA fluorescence in the NAc (**b**) or DLS (**d**) for the sessions marked by colored circles in **a** and **c**, which differ primarily in motivation rather than free consumption. **e**, **f**, Demand curves for the sessions marked by colored circles in **a** and **c**. The demand curves have similar y-intercepts but differ in slope.

**Extended Data Fig. 4:**
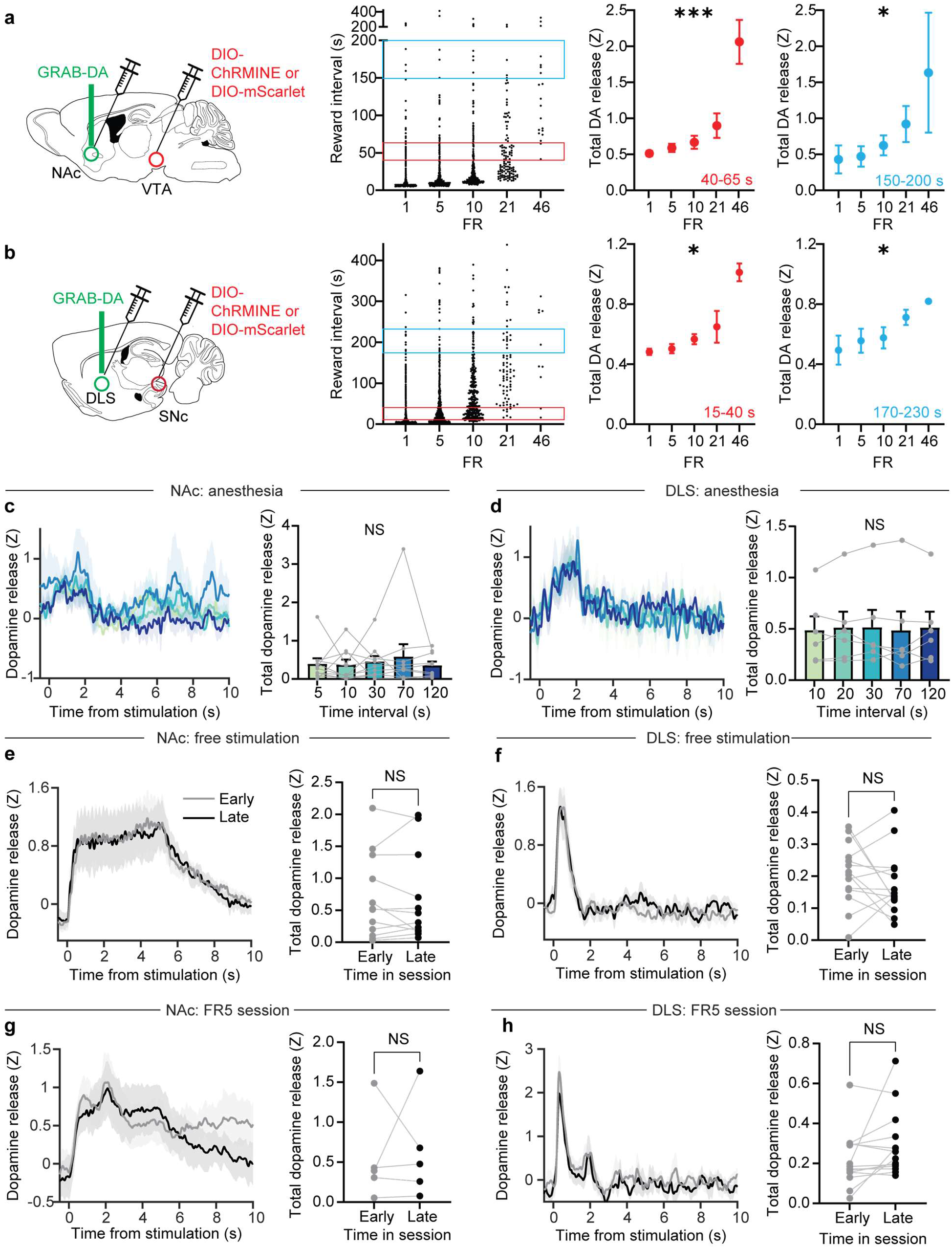
The effect of FR on optogenetically-evoked DA release does not depend on inter-stimulation interval or timing. **a, b,** Left, Distribution of inter-reward intervals as a function of FR for mice with fibers in the NAc (**a**) or DLS (**b**). Middle, Right, AUC of GRAB-DA photometry traces in response to optogenetic stimulation on trials with short (Middle, **a**: Kruskal-Wallis test, Kruskal-Wallis statistic=74.5, P<0.0001; n=2946 trials from 9 mice; **b**: Kruskal-Wallis test, Kruskal-Wallis statistic=10.8, P=0.029; n=1288 trials from 14 mice) or long (Right, **a**: Kruskal-Wallis test, Kruskal-Wallis statistic=10.1, P=0.04; n=25 trials from 9 mice; **b**: Kruskal-Wallis test, Kruskal-Wallis statistic=9.45, P=0.05; n=78 trials from 14 mice) inter-reward intervals. **c**, **d**, Left, Z-scored photometry traces of GRAB-DA fluorescence in the NAc (**c**) or DLS (**d**) in response to optogenetic stimulation at different intervals for mice under anesthesia. Right, AUC of Z-scored photometry traces in the NAc (**c**: Friedman test, Friedman statistic=3.28, P=0.51; n=10) or DLS (**d**: Friedman test, Friedman statistic=0.98, P=0.91; n=6) as a function of inter-stimulation interval. Intervals were chosen based on the average inter-reward intervals for each FR in the demand task. **e**, **f**, Left, Z-scored photometry traces of GRAB-DA fluorescence in the NAc (**e**) or DLS (**f**) during a session in which stimulation was given at random intervals without any nosepoking requirement. Right, AUC of Z-scored photometry traces in the NAc (**e**: Wilcoxon matched-pairs signed rank test, W=-6, P=0.85; n=12) or DLS (**f**: Wilcoxon matched-pairs signed rank test, W=-27, P=0.43; n=14) in the first half or last half of each free-reward session. **g**, **h**, Left, Z-scored photometry traces of GRAB-DA fluorescence in the NAc (**g**) or DLS (**h**) during a session in which stimulation was always given at FR5. Right, AUC of Z-scored photometry traces in the NAc (**g**: Wilcoxon matched-pairs signed rank test, W=1, P>0.99; n=5) or DLS (**h**: Wilcoxon matched-pairs signed rank test, W=61, P=0.058; n=14) in the first half or last half of each FR5 session. NS, not significant; *P<0.05, ***P<0.001. Error bars denote s.e.m.

**Extended Data Fig. 5:**
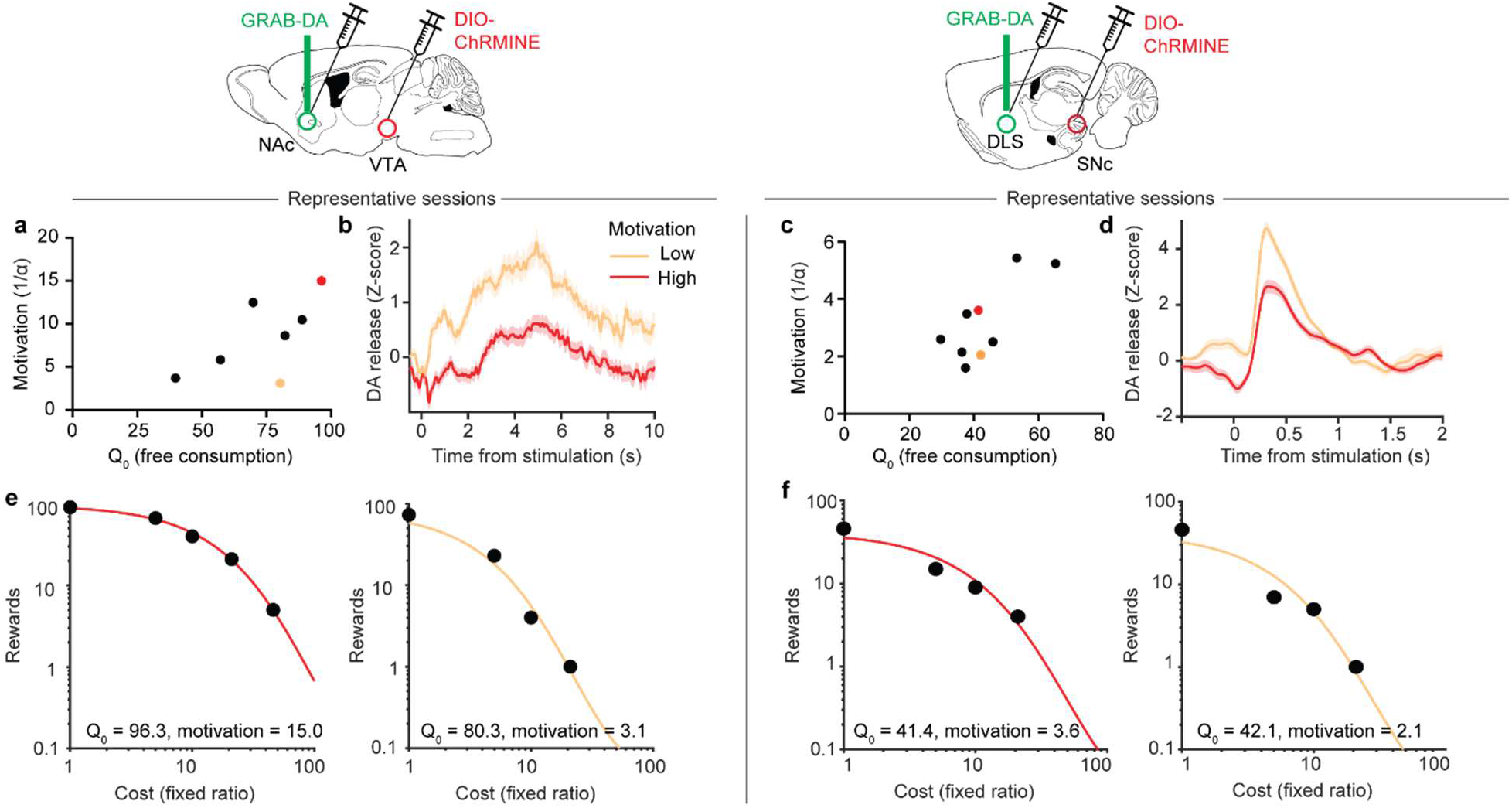
Representative examples of the inverse relationship between motivation and DA release for optogenetic rewards. **a**, **c**, Free consumption parameter versus motivation parameter extracted from individual sessions in one representative mouse with stimulation and recording in the NAc (**a**) or DLS (**c**). **b**, **d**, Z-scored photometry traces of stimulation-evoked GRAB-DA fluorescence in the NAc (**b**) or DLS (**d**) for the sessions marked by colored circles in **a** and **c**, which differ primarily in motivation rather than free consumption. **e**, **f**, Demand curves for the sessions marked by colored circles in **a** and **c**. The demand curves have similar y-intercepts but differ in slope.

